# Approximation of Indel Evolution by Differential Calculus of Finite State Automata

**DOI:** 10.1101/2020.06.29.178764

**Authors:** Ian Holmes

**Affiliations:** Department of Bioengineering, University of California, Berkeley

## Abstract

We introduce a systematic method of approximating finite-time transition probabilities for continuous-time insertion-deletion models on sequences. The method uses automata theory to describe the action of an infinitesimal evolutionary generator on a probability distribution over alignments, where both the generator and the alignment distribution can be represented by Pair Hidden Markov Models (Pair HMMs). In general, combining HMMs in this way induces a multiplication of their state spaces; to control this, we introduce a coarse-graining operation to keep the state space at a constant size. This leads naturally to ordinary differential equations for the evolution of the transition probabilities of the approximating Pair HMM. The TKF model emerges as an exact solution to these equations for the special case of single-residue indels. For the general case, the equations can be solved by numerical integration. Using simulated data we show that the resulting distribution over alignments, when compared to previous approximations, is a better fit over a broader range of parameters. We also propose a related approach to develop differential equations for sufficient statistics to estimate the underlying instantaneous indel rates by Expectation-Maximization. Our code and data are available at https://github.com/ihh/trajectory-likelihood.

## 1 Introduction

In molecular evolution, the equations of motion describe continuous-time Markov processes on discrete nucleotide or amino acid sequences. For substitution processes, these equations are reasonably well-understood, but insertion and deletions (indels) have proved elusive.

In models where a sequence is subject only to point substitutions at independently evolving sites, the likelihood can be factorized into a product of small Markov chains [39], solved exactly for an ancestor-descendant pair by considering the eigenstructure of the matrix exponential [41, 23], and extended to multiple aligned sequences by applying the sum-product algorithm to the phylogenetic tree [18]. These results are widely used in bioinformatics. Latent variables can be introduced to model rate heterogeneity [86] or selection [87], and the rate parameters estimated efficiently by Expectation-Maximization [33, 28]. The site-independent point substitution process can be generalized to the case where substitution rates are influenced by neighboring residues—or where substitution events simultaneously affect multiple residues—by expanding the matrix exponential as a Taylor series in neighboring contexts [48], extending to multiple alignments using variational (mean-field) approaches [37, 85].

By comparison, continuous-time Markov chain models of the indel process are tricky. The first—and only exactly-solved—example is the Thorne-Kishino-Felstenstein (TKF) model, which allows only single-residue indels. The TKF model reduces exactly to a linear birth-death process with immigration [78], which allows the joint distribution over ancestor-descendant sequence alignments to be expressed as a Hidden Markov Model (HMM) [32] that can be formally extended to multiple sequences using algebraic composition of automata [74, 25, 29, 83, 4]. This allows a statistical unification of alignment and phylogeny [47, 67, 76, 60, 61, 6, 81, 84, 1, 27, 34]. However, in practice, the TKF model itself is mostly used for inspiration in such applications, since its restriction to single-residue indel events is not consistent with empirical data [65, 9, 75, 8] and the consequent over-counting of events causes artefacts in statistical inference of alignments, trees, and rate parameters [79, 26, 32].

Attempts to generalize the TKF model to the more biologically-plausible case of multiple-residue indel events fall into two categories: those that attempt to analyze the process from first principles to arrive at finite-time transition probabilities [54, 43, 53, 17, 16, 15, 11], and those that guess closed-form approximations to these probabilities without such *ab initio* justifications [79, 56, 80, 68, 70, 69, 5]. In this paper we focus on the former type of approach. The latter approaches often proceed by breaking the sequence into indivisible multiple-residue fragments—or introducing other latent variables—but lacking any analytic connection of the fragment sizes or other newly-introduced parameters to the infinitesimal mutation rates of the underlying process, their evaluation in a statistical framework must necessarily be somewhat heuristic [11].

Formal mathematical treatment of the multi-residue indel process begins with Miklós and Toroczkai’s analysis of a model that allows long insertions but only single-residue deletions [54]. They developed a generating function for the gap length distribution, and used the method of characteristics to solve the associated partial differential equations. Arguably the most important feature of this model is that the alignment likelihood remains factorizable and associated with an HMM (albeit one with infinite states). This remains true for indel processes that allow both insertions and deletions to span multiple residues, under certain assumptions of spatial homogeneity [43, 53, 16], a theoretical result that helps to justify HMM-based approximations. However, calculating the transition probabilities of these HMMs from first principles is still nontrivial. Miklós *et al* [53], formalizing intuition of Knudsen and Miyamoto, [43], obtained reasonable approximations for short evolutionary time intervals by calculating exact likelihoods of short trajectories in the continuous-time Markov process. However, exhaustively enumerating these trajectories is extremely slow, and effectively impossible for trajectories with more than three overlapping indel events, so this approach is of limited use.

A recent breakthrough in this area was made by De Maio [11]. Starting from the approximation that the alignment likelihood can be factored into separate geometric distributions for insertion and deletion lengths, he derived ordinary differential equations (ODEs) for the evolution of the mean lengths of these distributions, yielding transition probabilities for the Pair HMM. De Maio’s method produces more accurate approximations to the multi-residue indel process than all previous attempts, though it has limitations: it’s restricted to models where the insertion and deletion rates are equal, does not (by design) include covariation between insertion and deletion lengths in the alignment, is inexact for the special case of the TKF model, and requires laborious manual derivation of the underlying ODEs.

In this paper, we build on De Maio’s results to develop a systematic differential calculus for finding HMM-based approxmate solutions of continuous-time Markov processes on strings which are “local” in the sense that the infinitesimal generator is an HMM. Our approach addresses the limitations of De Maio’s approach, identified in the previous paragraph. It does not require that insertion and deletion rates are equal, or that the process is time-reversible: any geometric distribution over indel lengths is allowed. It does account for covariation between insertion and deletion gap sizes. The TKF model emerges as a special case and the closed-form solutions to the TKF model are exact solutions to our model. Finally, although our equations can be derived without computational assistance, the analysis is greatly simplified by the use of symbolic algebra packages: both for the manipulation of equations, for which we used Mathematica [36], and for the manipulation of state machines, for which we used our recently published software Machine Boss [73].

The central idea of our approach is that the application of the infinitesimal generator to the approximating HMM generates a more complicated HMM that, by a suitable coarse-graining operation, can be mapped back to the simpler structure of the approximating HMM. By matching the expected transition usages of these HMMs, we derive ODEs for the transition probabilities of the approximator. Our approach is justified by improved results in simulations, yielding greater accuracy and generality than all previous approaches to this problem, including De Maio’s moment-based method (which can be seen as a version of our method that considers only indel-extending transitions in a symmetric model). Our approach is further justified by the emergence of the TKF model as an exact special case, without the need to introduce any additional latent variables such as fragment boundaries, or arbitrary time-dependence to the TKF formulae.

While we focus here on the multi-residue indel process, the generality of the infinitesimal automata suggests that other local evolutionary models, such as those allowing neighbor-dependent substitution and indel rates, might also be productively analyzed using this approach.

## 2 Methods

### 2.1 Expected transition usage

Suppose that we have a hidden Markov model, 𝕄, with *K* states which can be partitioned into match *σ*_*M*_, insert *σ*_*I*_, delete *σ*_*D*_, and null *σ*_*N*_ states, as in [14]. As a shorthand we will write *σ*_*ID*_ for *σ*_*I*_ ∪ *σ*_*D*_, and so on. Thus *σ*_*MIDN*_ is the complete set of *K* states.

We’ll be considering models with only one match state, which by convention will always be the first state, so *σ*_*M*_ = {1}.

Let *ϕ* = (*ϕ*_0_, *ϕ*_1_, … *ϕ*_*k*_) denote a state path.

The transition probability matrix is **Q** with elements *Q*_*ij*_ = *P* (*ϕ*_*k*+1_ = *j*|*ϕ*_*k*_ = *i*).

Let Φ(*X* → *Y* → *Z*) denote the set of state paths with the following properties:

- The path begins in a *σ*_*X*_ -state;
- The path ends in a *σ*_*Z*_ -state;
- For paths with more than two states, the intermediate states are all *σ*_*Y*_ -states.
- Let **J**^(*X*→*Y*)^ be a matrix that selects transitions from *σ*_*X*_ to *σ*_*Y*_

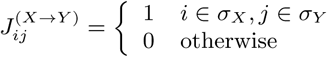

and let **Q**^(*X*→*Y*)^ *≡* **J**^(*X*→*Y*)^ ◦ **Q** where ◦ is the Hadamard product.

Consider a random path *ϕ* ∈ Φ(*M* → *IDN* → *M*) that begins and ends in the match state, passing only through non-match states in between. Let *T*_*XY*_ (*ϕ*) = |{(*i, j*) : *j* > *i*, (*ϕ*_*i*_ … *ϕ*_*j*_) ∈ Φ(*X* → *N* → *Y*)}| count the number of subpaths of *ϕ* that go from a state in *σ*_*X*_, via zero or more null states, to a state in *σ*_*Y*_. Thus if null states were removed from the path, then *T*_*XY*_ would be the number of transitions from *X*-states to *Y* -states.

The expectation of *T*_*XY*_ is

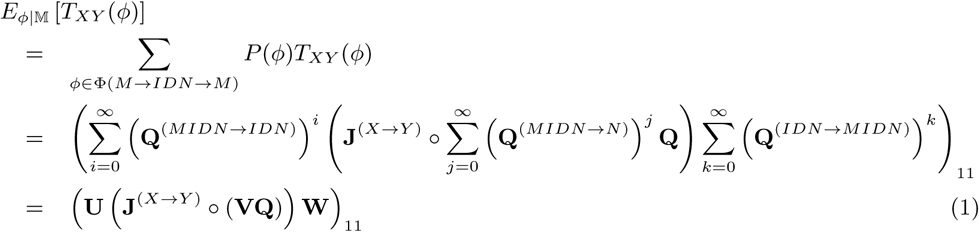

with

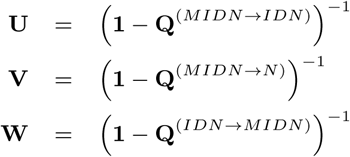

where 1 is the *K* × *K* identity matrix.

Let *S*_*X*_ (*ϕ*) be the number of *X*-states in *ϕ*, excluding the final state.Thus

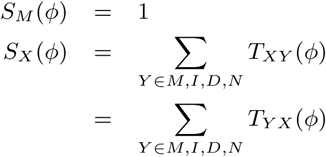

### 2.2 Three-state HMM

Consider the machine 𝔽(*t*) shown in Figure 1a with *σ*_*M*_ = {1}, *σ*_*I*_ = {2}, *σ*_*D*_ = {3}, and

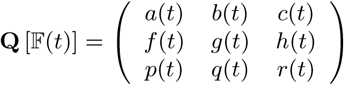

with *a* + *b* + *c* = 1, *f* + *g* + *h* = 1 and *p* + *q* + *r* = 1.

**Figure 1:**
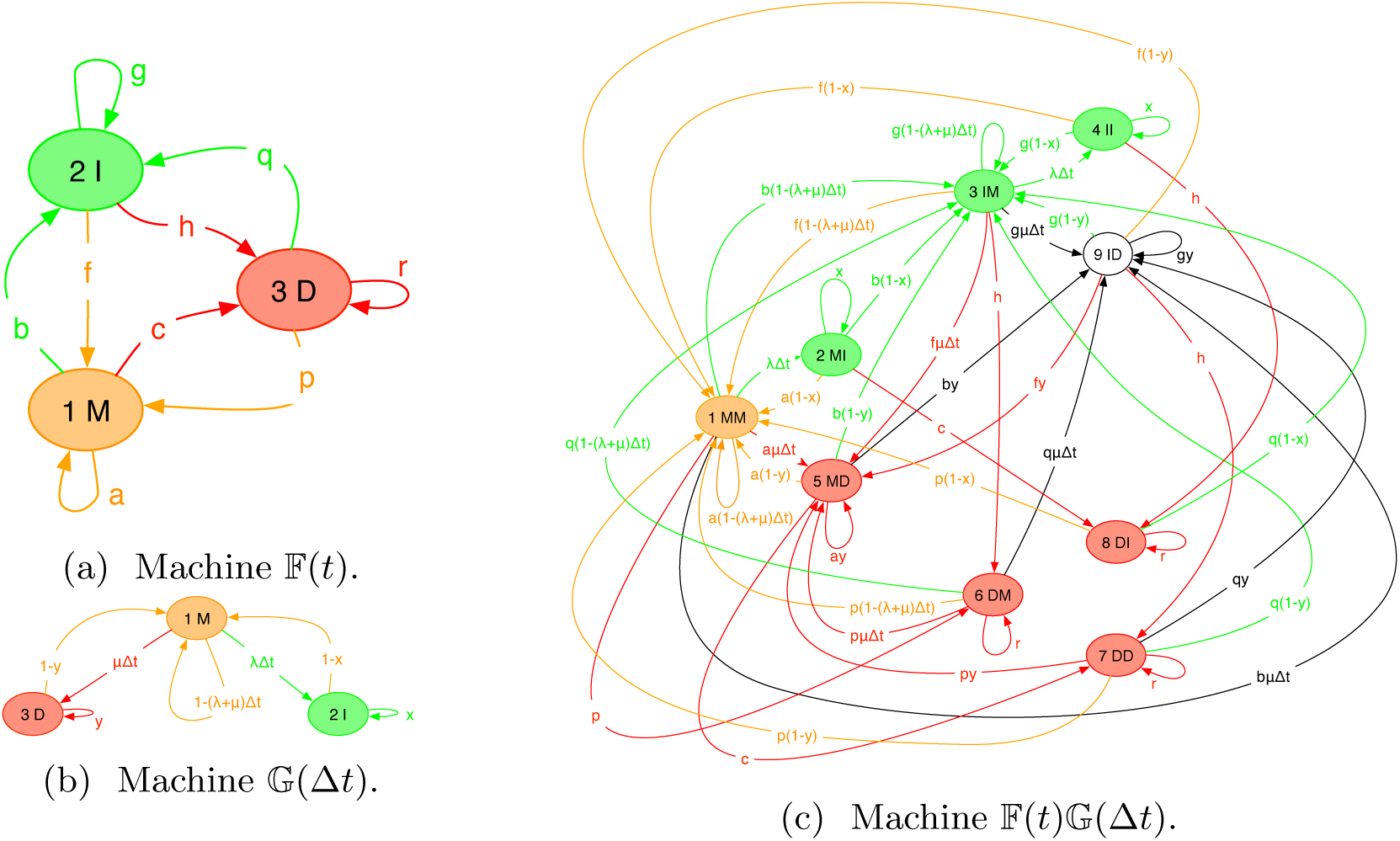
Three state machines for modeling indels in alignments. Match states are orange; insert states, green; delete states, red; and null states are uncolored. Transitions are colored by destination state. (1a) Machine 𝔽(*t*) models alignments at divergence time *t*. (1b) Machine 𝔾(Δ*t*) models the infinitesimal evolution over time Δ*t*. (1c) Machine 𝔽(*t*)𝔾(Δ*t*), the transducer product of the previous two machines, models alignments at divergence time *t* + Δ*t*. Our approach is to approximate this with a machine of the same form as (1a).

Here *t* will play the role of a time parameter. Let

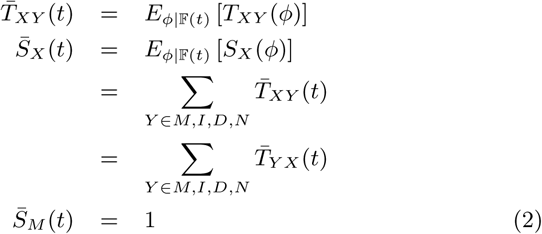

where the expectations are as defined in (1). (Throughout this paper, such expectations are over *ϕ* ∈ Φ(*M* → *IDN* → *M*).) Evidently,

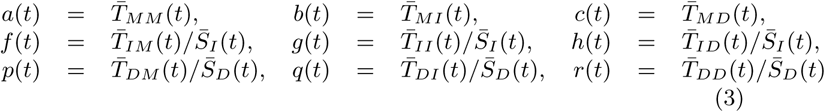

By (1),

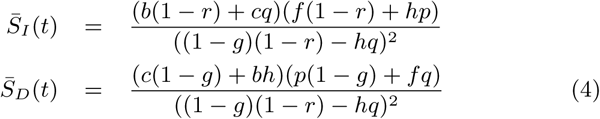

The essence of our approach is to use transducer composition to study infinitesimal increments in 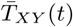 and thereby obtain differential equations that can be solved to find these parameters.

### 2.3 Rate of change of expected transition counts

The infinitesimal transducer 𝔾(Δ*t*) of Figure 1b has states *σ*_*M*_ = {1}, *σ*_*I*_ = {2}, *σ*_*D*_ = {3}, and transition matrix

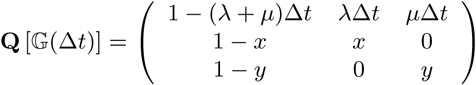

This describes a spatially homogeneous “long indel” model where the indel lengths are geometrically distributed. The model parameters are the insertion and deletion rates (*λ, µ*) and extension probabilities (*x, y*), and the time interval Δ*t* ≪ 1.

Composing 𝔽(*t*) (Figure 1a) with 𝔾(Δ*t*) (Figure 1b) yields 𝔽(*t*)𝔾(Δ*t*), the machine of Figure 1c. This machine^1^ has states *σ*_*M*_ = {1}, *σ*_*I*_ = {2, 3, 4}, *σ*_*D*_ = {5, 6, 7, 8}, *σ*_*N*_ = {9}, and transition matrix

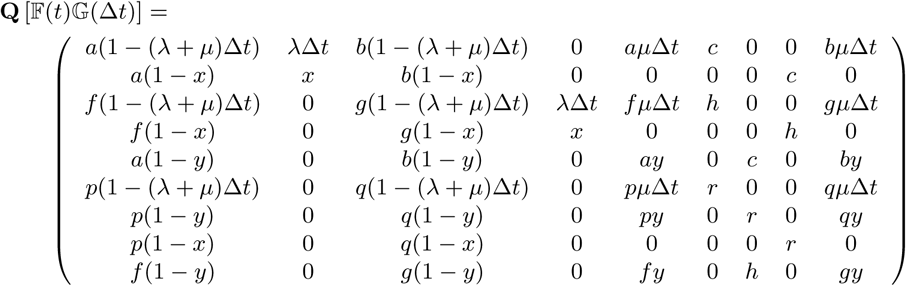

We now make the approximation 𝔽(*t*)𝔾(Δ*t*) ≈ 𝔽(*t* + Δ*t*) which is to say that the 9-state machine of Figure 1c can be approximated by the 3-state machine of Figure 1a by infinitesimally increasing the time parameter of the simpler machine. This will not, in general, be exact (with the exception of the TKF model, discussed in Section 2.4). However, by mapping states (and hence transitions) of 𝔽𝔾 back to 𝔽, and setting 𝔽’s transition probabilities proportional to the expected number of times these transitions are used in 𝔽𝔾, we find a maximum-likelihood fit.

The expected transition counts evolve via the coupled differential equations

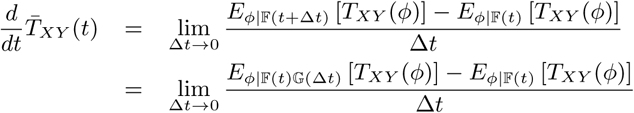

Expanding (1) to first order in Δ*t* and then taking the limit Δ*t* → 0, we arrive at the following equations for the expected transition counts^2^

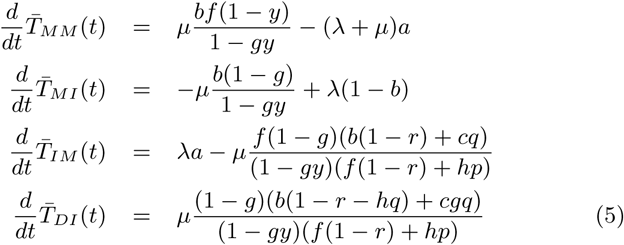

with boundary condition

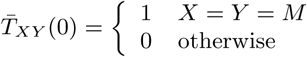

The parameters (*a, b, c, f, g, h, p, q, r*) are defined by (3) for *t* > 0, but must also be specified as part of the boundary condition at *t* = 0, where *a*(0) = *f* (0) = *p*(0) = 1 and *b*(0) = *c*(0) = *g*(0) = *h*(0) = *q*(0) = *r*(0) = 0.

The remaining counts are obtained from (2)

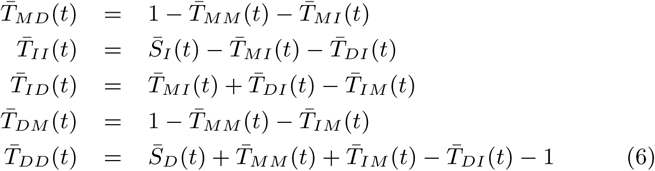

The expected state occupancies are governed by the following equations

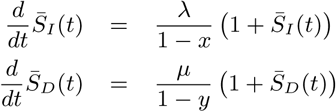

which, generalizing [11], have the closed-form solution

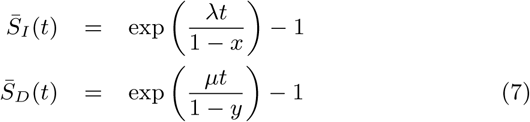

### 2.4 TKF model

When *x* = *y* = 0, our model reduces to the TKF model [78]. It is known [32] that the solution to the TKF model can be expressed as a Pair HMM of the form shown in Figure 1a, with parameters

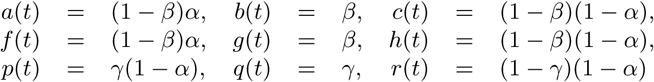

Where

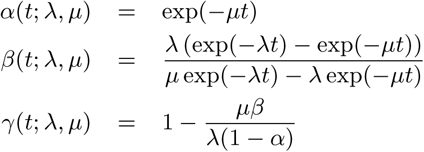

It can readily be verified that this is an exact solution to equations (3) through (7) when *x* = *y* = 0. Thus, our model reduces exactly to the TKF model when indels involve only single residues. In this case, the equivalence 𝔽(*t* + Δ*t*) = 𝔽(*t*)𝔾(Δ*t*) is exact in the limit Δ*t* → 0.

### 2.5 Parameterization via EM

Sufficient statistics for parameterizing the infinitesimal process of Figure 1b are

- *S*, the number of sequences in the dataset;
- *n*.^*l*^, the number of sites at which deletions can occur, integrated over time (.*l/t* is the mean sequence length over the time interval);
- *n*^*λ*^, the number of insertion events that occurred;
- *n*^*µ*^, the number of deletion events that occurred;
- *n*^*x*^, the number of insertion extensions (*n*^*x*^+ 1 is the total number of inserted residues);
- *n*^*y*^, the number of deletion extensions (*n*^*y*^ + 1 is the total number of deleted residues).

Given these statistics, the maximum likelihood parameterization is^3^

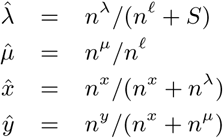

Let 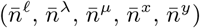 denote the expectations of the sufficient statistics over the posterior distribution of histories. The Expectation Maximization algorithm for continuous-time Markov processes alternates between calculating these posterior expectations for some parameterization (*λ*_*k*_, *µ*_*k*_, *x*_*k*_, *y*_*k*_) and using them to find a better parameterization [33, 28, 31, 13]

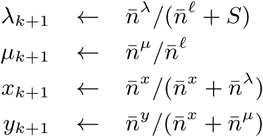

In any state path through the machines of Figure 1, each transition will make an additive contribution to these statistics. Let 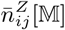 denote the contribution to 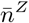 made by transition *i* → *j* of machine 𝕄. With reference to Figure 1c, and by the rules of algebraic automata composition [83], each state of 𝔽(*t*)𝔾(Δ*t*) can be written as a tuple (*i, j*) of a 𝔽-state *i* and a 𝔾-state *j*, and each transition weight of 𝔽(*t*)𝔾(Δ*t*) takes the form

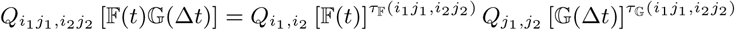

where *τ*_𝕄_(*i*_1_*j*_1_, *i*_2_*j*_2_) is 1 if machine 𝕄 changes state in the transition (*i*_1_, *j*_1_) → (*i*_2_, *j*_2_), and 0 if it does not. Using this, we can write

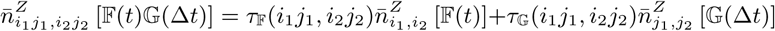

Where

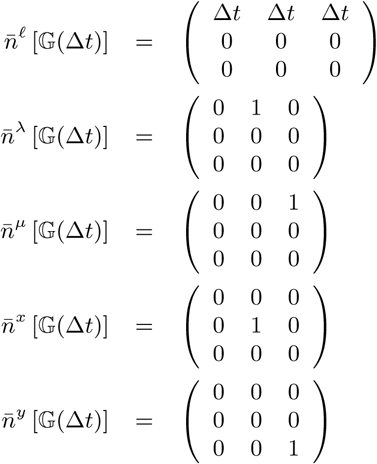

and thus, for 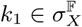 and 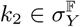,

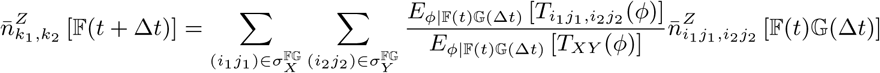

where *E*[*T*_*ij*_] is defined by (1) in the same way for transitions between individual states as *E*[*T*_*XY*_] is defined for transitions between sets of states (e.g. by defining *σ*_*i*_ *≡* {*i*} for *i* ∈ {1 … *K*}). **Conjecture**. Expanding these equations to first order in Δ*t* and taking the limit Δ*t* → 0 leads to ODEs for 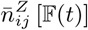, analogous to the ODEs for 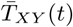 given in (5), that can be used to fit the parameters of the infinitesimal generator from unaligned sequence data by weighting the 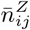 with the posterior transition usage counts obtained using the Baum-Welch algorithm.

### 2.6 Higher moments

We here include a few results relating to our model that may be useful, but are not directly needed to derive the differential equations that govern it.

The matrix method of Section 2.1 can be used to find *E*[*S*_*I*_] and *E*[*S*_*D*_] directly, as well as higher moments. Let **X** ∈ {**I, D**} be the diagonal matrix indicating membership of *σ*_*X*_, so *X*_*ij*_ = *δ*(*i* = *j*)*δ*(*i* ∈ *σ*_*X*_). Then

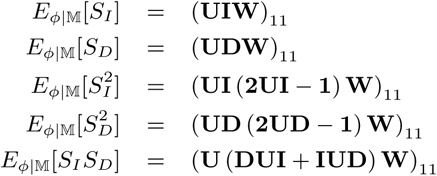

In the case of machine 𝔽 in Figure 1a, the first two of these moments have already been given in (4). The others are

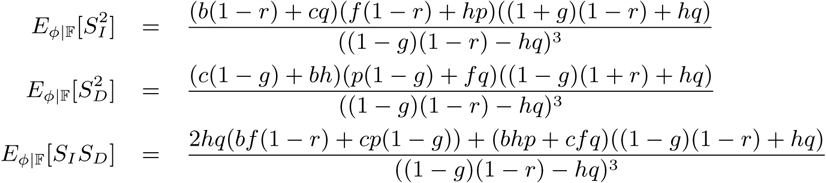

These results can also be obtained from the moment generating function for the joint distribution *P* (*S*_*I*_, *S*_*D*_ |𝔽), which is

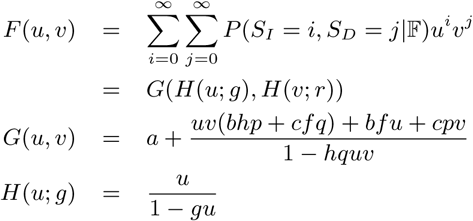

where *G* is the generating function for *P* (*T*_→*I*_, *T*_→*D*_ |𝔽) where *T*_→*I*_ = *T*_*MI*_ + *T*_*DI*_ and *T*_→*D*_ = *T*_*MD*_ + *T*_*ID*_, and *H* is the generating function for a geometric series.

## 3 Results

We implemented the likelihood calculations of Section 2.3 and those of De Maio (2020) and Miklós, Lunter and Holmes (2004), which are directly comparable, although De Maio requires that *λ* = *µ* and the method of Miklós *et al* is only practical for high-throughput evaluation up to gap lengths of about 30 (Figure 2). As a shorthand, we refer to these methods as DM20 [11], MLH04 [53], and H20 for the current method. We also implemented a simulator for the underlying indel process and compared summary statistics across the different methods, including the relative entropy of the joint distribution *P* (*S*_*I*_, *S*_*D*_) (although for MLH04 it was necessary to renormalize this to *P* (*S*_*I*_, *S*_*D*_ |*S*_*I*_ ≤ 30, *S*_*D*_ ≤ 30) to avoid infinities in the relative entropy) and various marginals and moments of this distribution. Our implementations and simulation results are available at https://github.com/ihh/trajectory-likelihood.

**Figure 2:**
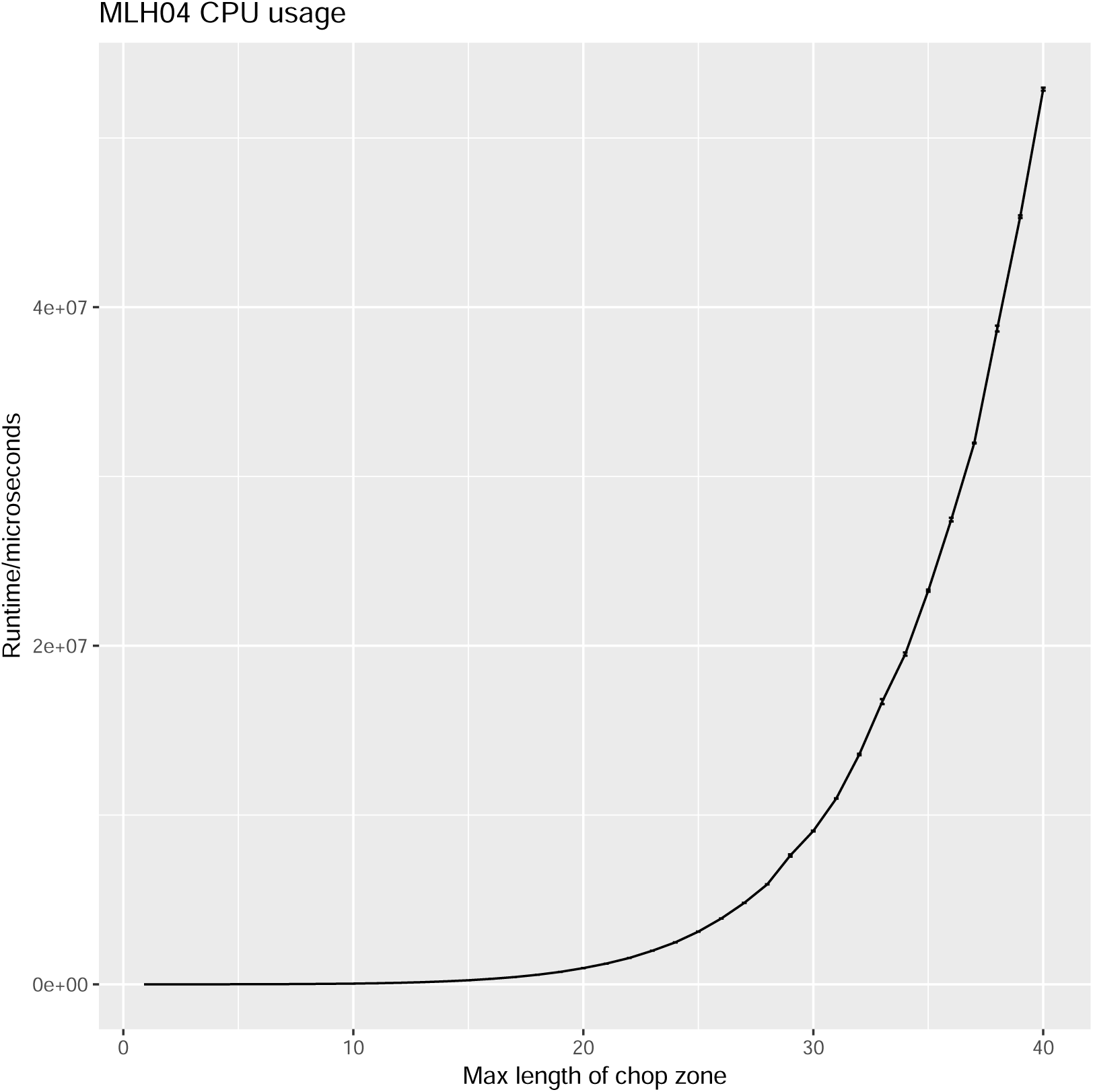
The running time of the MLH04 method is prohibitive and increases steeply for long gaps, since it requires the explicit enumeration of all intermediate indel states in the trajectory [53]. In this plot, each datapoint is an average of 100-1000 repetitions on a late-2014 iMac (4GHz quad-core Intel i7 CPU, 32GB 1600MHz DDR3 RAM). The running times of DM20 and MLH04 are negligible in comparison (< 1ms).

We performed simulations at parameter settings *µ* ∈ {0.05, 0.5}, *λ* ∈ {*µ*, 0.9*µ*}, *y* ∈ {0.5, 0.75, 0.9}, *x* ∈ {*λy/µ*, 0.45, 0.8, 0.95}, and *t* ∈ {2^*k*^ :−6 ≤ *k* ≤ 3}. Note that the existence of a reversible equilibrium by detailed balance requires *λ* < *µ* [78] and *µx* = *λy* [53]. When *x* is larger than this, the sequence length tends to increase without limit, bringing simulations to a halt. The experiments with *x* > *λy/µ* were abandoned at higher values of *t* for this reason.

In each experiment we simulated the random indel drift of a 1kb sequence. For *t* > 1, we performed 10^5^ repetitions of each simulation experiment; for *t* ≤ 1, we performed 10^6^ repetitions.

Summary statistics for some of these experiments are plotted in Figure 3 and some relative entropies are also given in Table 1. In almost all cases, the new method is a closer approximation than DM20. The MLH04 trajectory-enumerating approximation is often a closer fit at short times (with the proviso that it can only handle short gaps, and takes significantly longer to compute) but it quickly fails when the number of indel events in a typical trajectory exceeds the 3 events that are computationally tractable to enumerate.

**Table 1:** Selected relative entropies of simulation benchmark results. In each row the best method is highlighted dark red, and the second-best is highlighted light red. In all cases shown, the insertion and deletion rates were equal (*λ* = *µ*). The “Rev?” column indicates whether a given parameter setting is time-reversible (*λy* = *µx*).

| | | | | | $D(P(S_I, S_D\vert \text{simulated})\vert \vert P(S_I, S_D\vert \text{approx}))$ | | |
| --- | --- | --- | --- | --- | --- | --- | --- |
| $\mu$ | $x$ | $y$ | $t$ | Rev? | DM20 [11] | MLH04 [53] | H20 |
| 0.05 | 0.5 | 0.45 | 0.125 | | $2.04985 \times 10^{-5}$ | $9.11216 \times 10^{-7}$ | $8.81290 \times 10^{-7}$ |
| 0.05 | 0.5 | 0.45 | 1 | | $1.68563 \times 10^{-4}$ | $2.69044 \times 10^{-6}$ | $1.69689 \times 10^{-5}$ |
| 0.05 | 0.5 | 0.45 | 8 | | $5.79716 \times 10^{-3}$ | $2.23302 \times 10^{-2}$ | $3.45910 \times 10^{-3}$ |
| 0.05 | 0.5 | 0.5 | 0.125 | ✓ | $1.22097 \times 10^{-6}$ | $1.17394 \times 10^{-6}$ | $1.22242 \times 10^{-6}$ |
| 0.05 | 0.5 | 0.5 | 1 | ✓ | $2.18927 \times 10^{-5}$ | $3.27733 \times 10^{-6}$ | $2.12916 \times 10^{-5}$ |
| 0.05 | 0.5 | 0.5 | 8 | ✓ | $5.42222 \times 10^{-3}$ | $3.00300 \times 10^{-2}$ | $4.20105 \times 10^{-3}$ |
| 0.05 | 0.5 | 0.8 | 0.125 | | $3.57032 \times 10^{-3}$ | $2.40144 \times 10^{-6}$ | $4.91577 \times 10^{-6}$ |
| 0.05 | 0.75 | 0.45 | 0.125 | | $4.26426 \times 10^{-4}$ | $9.50348 \times 10^{-7}$ | $1.11389 \times 10^{-6}$ |
| 0.05 | 0.75 | 0.45 | 1 | | $3.38901 \times 10^{-3}$ | $2.94778 \times 10^{-6}$ | $8.41352 \times 10^{-6}$ |
| 0.05 | 0.75 | 0.45 | 8 | | $2.93007 \times 10^{-2}$ | $1.31450 \times 10^{-3}$ | $1.73884 \times 10^{-3}$ |
| 0.05 | 0.75 | 0.75 | 0.125 | ✓ | $3.03606 \times 10^{-6}$ | $2.86492 \times 10^{-6}$ | $3.03568 \times 10^{-6}$ |
| 0.05 | 0.75 | 0.75 | 1 | ✓ | $1.83832 \times 10^{-5}$ | $8.69690 \times 10^{-6}$ | $1.81854 \times 10^{-5}$ |
| 0.05 | 0.75 | 0.75 | 8 | ✓ | $3.75601 \times 10^{-3}$ | $9.11449 \times 10^{-3}$ | $3.32154 \times 10^{-3}$ |
| 0.05 | 0.75 | 0.8 | 0.125 | | $6.93445 \times 10^{-5}$ | $2.84201 \times 10^{-6}$ | $3.27834 \times 10^{-6}$ |
| 0.05 | 0.9 | 0.45 | 0.125 | | $4.59847 \times 10^{-4}$ | $7.05138 \times 10^{-7}$ | $1.28355 \times 10^{-6}$ |
| 0.05 | 0.9 | 0.45 | 1 | | $3.68176 \times 10^{-3}$ | $2.30351 \times 10^{-6}$ | $5.26999 \times 10^{-6}$ |
| 0.05 | 0.9 | 0.45 | 8 | | $3.01802 \times 10^{-2}$ | $1.62709 \times 10^{-5}$ | $6.49049 \times 10^{-4}$ |
| 0.05 | 0.9 | 0.8 | 0.125 | | $1.40027 \times 10^{-4}$ | $2.11248 \times 10^{-6}$ | $2.84768 \times 10^{-6}$ |
| 0.05 | 0.9 | 0.9 | 0.125 | ✓ | $5.83560 \times 10^{-6}$ | $4.19581 \times 10^{-6}$ | $5.83597 \times 10^{-6}$ |
| 0.05 | 0.9 | 0.9 | 1 | ✓ | $2.94696 \times 10^{-5}$ | $4.60033 \times 10^{-5}$ | $2.94067 \times 10^{-5}$ |
| 0.05 | 0.9 | 0.9 | 8 | ✓ | $1.59644 \times 10^{-3}$ | $2.88432 \times 10^{-3}$ | $1.45752 \times 10^{-3}$ |
| 0.5 | 0.5 | 0.45 | 0.125 | | $2.22362 \times 10^{-4}$ | $4.17791 \times 10^{-6}$ | $3.15751 \times 10^{-5}$ |
| 0.5 | 0.5 | 0.45 | 1 | | $9.41950 \times 10^{-3}$ | $5.63014 \times 10^{-2}$ | $5.59983 \times 10^{-3}$ |
| 0.5 | 0.5 | 0.45 | 8 | | $6.20265 \times 10^{-1}$ | $1.36809\text{e}+01$ | $8.83542 \times 10^{-2}$ |
| 0.5 | 0.5 | 0.5 | 0.125 | ✓ | $3.82629 \times 10^{-5}$ | $3.70476 \times 10^{-6}$ | $3.68489 \times 10^{-5}$ |
| 0.5 | 0.5 | 0.5 | 1 | ✓ | $9.37828 \times 10^{-3}$ | $7.47712 \times 10^{-2}$ | $6.79378 \times 10^{-3}$ |
| 0.5 | 0.5 | 0.5 | 8 | ✓ | $5.16437 \times 10^{-1}$ | $1.37034\text{e}+01$ | $9.75359 \times 10^{-2}$ |
| 0.5 | 0.5 | 0.8 | 0.125 | | $3.78591 \times 10^{-2}$ | $5.03568 \times 10^{-4}$ | $1.41159 \times 10^{-3}$ |
| 0.5 | 0.75 | 0.45 | 0.125 | | $4.24415 \times 10^{-3}$ | $1.89067 \times 10^{-6}$ | $1.16934 \times 10^{-5}$ |
| 0.5 | 0.75 | 0.45 | 1 | | $3.77702 \times 10^{-2}$ | $3.42854 \times 10^{-3}$ | $3.13205 \times 10^{-3}$ |
| 0.5 | 0.75 | 0.45 | 8 | | $8.54601 \times 10^{-1}$ | $9.90949 \times 10^{-1}$ | $2.84669 \times 10^{-1}$ |
| 0.5 | 0.75 | 0.75 | 0.125 | ✓ | $2.17573 \times 10^{-5}$ | $5.14187 \times 10^{-6}$ | $2.13362 \times 10^{-5}$ |
| 0.5 | 0.75 | 0.75 | 1 | ✓ | $6.80341 \times 10^{-3}$ | $2.20785 \times 10^{-2}$ | $5.83371 \times 10^{-3}$ |
| 0.5 | 0.75 | 0.75 | 8 | ✓ | $6.25327 \times 10^{-1}$ | $2.45203\text{e}+00$ | $3.02844 \times 10^{-1}$ |
| 0.5 | 0.75 | 0.8 | 0.125 | | $6.99602 \times 10^{-4}$ | $2.01321 \times 10^{-5}$ | $4.86974 \times 10^{-5}$ |
| 0.5 | 0.9 | 0.45 | 0.125 | | $4.59746 \times 10^{-3}$ | $2.79371 \times 10^{-6}$ | $8.60162 \times 10^{-6}$ |
| 0.5 | 0.9 | 0.45 | 1 | | $3.80985 \times 10^{-2}$ | $2.94801 \times 10^{-5}$ | $1.13099 \times 10^{-3}$ |
| 0.5 | 0.9 | 0.45 | 8 | | $4.67302 \times 10^{-1}$ | $1.44795 \times 10^{-2}$ | $1.96366 \times 10^{-1}$ |
| 0.5 | 0.9 | 0.8 | 0.125 | | $1.40555 \times 10^{-3}$ | $1.38761 \times 10^{-5}$ | $1.65939 \times 10^{-5}$ |
| 0.5 | 0.9 | 0.9 | 0.125 | ✓ | $3.65783 \times 10^{-5}$ | $6.86812 \times 10^{-5}$ | $3.64414 \times 10^{-5}$ |
| 0.5 | 0.9 | 0.9 | 1 | ✓ | $2.86092 \times 10^{-3}$ | $4.97623 \times 10^{-3}$ | $2.55231 \times 10^{-3}$ |
| 0.5 | 0.9 | 0.9 | 8 | ✓ | $4.92217 \times 10^{-1}$ | $2.11573 \times 10^{-1}$ | $3.28780 \times 10^{-1}$ |

**Figure 3:**
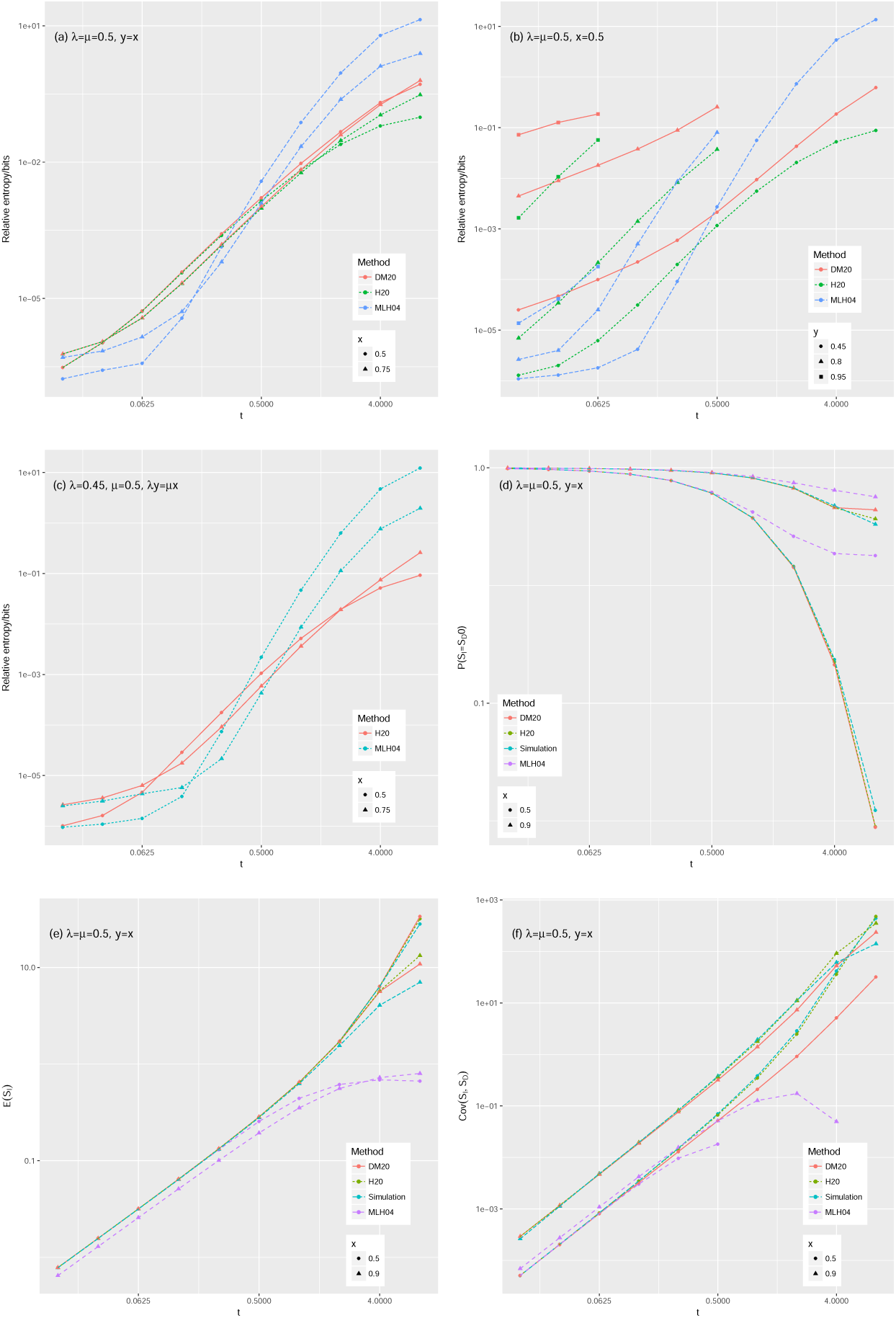
Summary statistics of simulation benchmarks plotted against evolutionary time. Shown are the relative entropies from the simulation distribution to the various approximations for reversible (a), irreversible-with-equal-rates (b), and reversible-with-inequal-rates (c) parameter regimes; the probability of observing no gap (d); the expected number of insertions (e); and the covariance between the number of insertions and deletions (f).

Despite its built-in assumption that insertion and deletion length distributions are separable (the random lengths are conditionally independent given that they are nonzero), DM20 does not perform too much worse than MLH04 at predicting covariation between these lengths (Figure 3f), perhaps because the greatest contribution to the covariation arises from the joint probability *P* (*S*_*I*_ = *S*_*D*_ = 0), which DM20 does explicitly model (Figure 3d).

We mostly report simulation results with *λ* = *µ* to allow comparison with DM20, though we did perform some simulations with *λ* = 0.9*µ* in order to compare H20 to MLH04 in this asymmetric-but-reversible regime. We note that the performance of H20 is not significantly worse when *λ* ≠ *µ* (Figure 3c). By contrast, when *λ* = *µ* but *x* > *y*, so the model is symmetric with respect to insertion/deletion rates but irreversible with respect to their lengths, both H20 and DM20 deteriorate; but DM20 more so than H20 (Figure 3b).

## 4 Discussion

We have shown that an evolutionary model which can be represented infinitesimally as an HMM can be formally connected to a Pair HMM that approximates its finite-time solution. This may be viewed as an automata-theoretic framing of the Chapman-Kolmogorov equation.

Ours is fundamentally a coarse-graining approach. It is generally the case that composing two state machines will yield a more complex machine, since the composite state space is the Cartesian product of the components. Our approach is to approximate this more complex machine with a renormalizing operation that eliminates null states and maps the remaining states of each type back to a single representative state in the approximator.

We have used this approach to derive ordinary differential equations for the transition probabilities of a minimal (three-state) Pair HMM that approximates a continuous-time indel process with geometrically distributed indel lengths. We have implemented numerical solutions to these equations, and demonstrated that they outperform the previous best methods. We have also conjectured similar ODEs for the posterior expectations of the sufficient statistics that would be required to fit this model by Expectation Maximization, though we have not yet tested this approach.

Point substitution models are the foundation of likelihood phylogenetics [35, 19]. There is, additionally, a substantial literature combining such models with HMMs [86, 20, 21, 45, 62, 72, 49, 24, 59, 12] and stochastic context-free grammars (SCFGs) [42, 63, 82, 77] for purposes of sequence annotation. The development of indel models has been slower. This may be, in large part, because integrating alignment and phylogeny is technically and computationally demanding. Multiple sequence alignments are a nuisance variable whose point estimation is a tolerable compromise when considering substitution processes, although several studies report that bias due to alignment error is a significant problem in substitution-founded phylogenetics [22, 77, 44, 50, 3] that must be handled with great care to avoid biasing inference [38, 64, 71]. When it comes to indel-based analysis, this compromise of conditioning on a single alignment rarely remains tenable, except perhaps in the “big data” limit, e.g. for closely-related sequences at genome scale [46, 66]. So indel-based phylogenetic inference must often co-sample or otherwise marginalize alignments, which is inherently harder [76, 60, 81, 34]. Nevertheless, the inexactitude of existing long-indel approximations may also have been a contributing obstacle to their slow adoption in the bioinformatic tool chain. If so, then the results presented here might help.

It seems possible that our method can be applied to other instantaneous rate models of local evolution where the infinitesimal generator can be represented as an HMM. It is tempting to speculate that a similar approach may also be productively applied to SCFGs [30, 7]. Such an approach would be more challenging; for example, elimination of null states from SCFGs is more complicated than for HMMs. One motivating goal would be to describe a realistic evolutionary drift process over RNA structures, with the goal of reconstructing the RNA world [52]. It’s also conceivable that approaches similar to those described here for biological sequences could be used to analyze phonemes [6], literary texts [2], music [10], source code [55], bird songs [40], or other alignable sequences that evolve over time.

## Footnotes

1 The transducer composition 𝔽(*t*) × 𝔾(Δ*t*) was performed using the automata algebra program Machine Boss [73]. A general procedure for doing this for any two machines 𝔸, 𝔹 involves taking the Cartesian product of the two machines’ state spaces and then synchronizing their transitions. This ensures that, if 𝕄_*XY*_ represents the result of the Forward algorithm for machine 𝕄 (with null states eliminated) and sequences *X, Y*, then (𝔸𝔹)_*XZ*_ = ∑ _*Y*_ 𝔸_*XY*_ 𝔹_*Y Z*_. More details on these operations can be found in [73, 83, 84] and their information-theoretic and linguistic roots in [51, 58, 57].

2 These calculations were performed using the symbolic algebra program Mathematica [36].

3 The formula for *λ* assumes that insertions can occur at the start and end of the sequence, as is usual [53]. Strictly, this requires that we specify a start and end state for 𝔾, rather than implicitly assuming infinite-length sequences as we have done up to this point. Specifically we start 𝔾 in the match state, and add transitions to the end state from the insert state with weight 1 − *x*, from the delete state with weight 1 − *y*, and from the match state with weight 1. This can be extended with rigor throughout the analysis by also specifying start and end states for 𝔽 and deriving differential equations for the transitions involving these states. Since it complicates the presentation to do this, we have omitted it. A heuristic for 𝔽 that is probably acceptable for most applications is to start it in the match state, and to allow transitions to the end state from the insert state with weight 1 − *g*, from the delete state with weight 1 − *q*, and from the match state with weight 1 − *b*.

